# Searching the optimal folding routes of a Complex Lasso protein

**DOI:** 10.1101/507079

**Authors:** Claudio Perego, Raffaello Potestio

## Abstract

Understanding how polypeptides can efficiently and reproducibly attain a self-entangled conformation is a compelling biophysical challenge, which might shed new light on our general knowledge of protein folding. Complex Lassos, namely self-entangled protein structures characterized by a covalent loop sealed by a cysteine bridge, represent an ideal test system in the framework of entangled folding. Indeed, as cysteine bridges form in oxidizing conditions, they can be used as on/off switches of the structure topology, to investigate the role played by the backbone entanglement in the process.

In the present work we have used molecular dynamics to simulate the folding of a complex lasso glycoprotein, Granulocyte-macrophage colony-stimulating factor, modeling both reducing and oxidizing conditions. Together with a well-established Go-like description, we have employed the elastic folder model, a Coarse-Grained, minimalistic representation of the polypeptide chain, driven by a structure-based angular potential. The purpose of this study is to assess the kinetically optimal pathways, in relation to the formation of the native topology. To this end we have implemented an evolutionary strategy that tunes the elastic folder model potentials to maximize the folding probability within the early stages of the dynamics. The resulting protein model is capable of folding with high success rate, avoiding the kinetic traps that hamper the efficient folding in the other tested models. Employing specifically designed topological descriptors, we could observe that the selected folding routes avoid the topological bottleneck by locking the cysteine bridge after the topology is formed.

These results provide valuable insights on the selection of mechanisms in self-entangled protein folding while, at the same time, the proposed methodology can complement the usage of established minimalistic models, and draw useful guidelines for more detailed simulations.

## INTRODUCTION

Almost a quarter of a century of research has been dedicated to the study of proteins that exhibit a self-entangled native fold. Nowadays, up to the 6% of the structures deposited in the Protein Data Bank (PDB)(1) are self-entangled proteins(2, 3). Since the first natively knotted protein was discovered in 1994(4), the existence of such topologically complex folds has represented a new challenge in the understanding of protein folding, fostering a wide range of studies. A number of reviews addressing the topic of self-entangled proteins can be found in the literature (see e.g. (2, 5–7), just to name the most recent), each addressing a different aspect of this variegated research field. The discovery of self-entangled protein structures has raised a few crucial questions related to their scarcity(8, 9), their conservation along evolution(10, 11), and their possible biological function(2, 7, 12–14).

In the present work we address the following question: how can the amino acid chain fold reproducibly and efficiently into a specific, nontrivial topology? Many experiments were conducted to answer this question, showing e.g. that these proteins can spontaneously tie themselves to the native topology(15); that the formation of the entanglement is a rate limiting step(15–17); and that one or few folding routes happen to be dominant, presumably representing the most efficient and reliable mechanisms(18). These crucial results demonstrate that self-entangled folding clearly differentiates from the simple picture of two-state folding of small, non-entangled proteins, but it is evident that efforts are still needed to reach a comprehensive and sound picture of this phenomenon.

In this framework, an interesting class of proteins is constituted by Complex Lassos (CLs)(19), entangled structures that exhibit a covalent loop closed by a disulphide bridge. The surface of this loop is pierced one or more times by the polypeptide chain, forming a non-trivial topology. Since Leptin was classified as the first CL protein(20), this topological state has been found to be widespread in the known PDB structures, characterizing about 18% of the proteins containing a cysteine bridge(21). Most of the CLs are secreted proteins, with signaling functions, and their topology is believed to have a crucial role in their biological activity(22, 23). Moreover, the topology of CLs can be controlled externally, since the cysteine bridge is stable in an oxidizing solution, while it does not form in a reducing environment. This feature allows one to directly study the effect of the topological barrier on the folding mechanism, electing CLs as ideal test systems for a deeper understanding of entangled folding.

As for simple proteins, the experimental probe of folding pathways in self-entangled proteins such as CLs can only provide indirect indications. For this reason Molecular Dynamics (MD) simulation represents an essential, complementary tool for the study of the process. We must however stress that the time duration of self-entangled folding typically exceeds the range accessible by all-atom simulations employing realistic interactions. This is the reason why, except in two notable cases(24, 25), the available computational results have been obtained using simplified, minimalistic protein models, which allow for a thorough sampling of the conformational space, while at the same time providing indications on the theoretical principles of the folding.

By far the most used methods are the so-called Gō models (GōM) (see e.g. (26, 27)), named after the pioneering work of Gō (28). In GōMs the protein is described as an hetero-polymer chain that encodes its native fold in the interaction potential. This kind of description stems out from the established Energy Landscape Theory (ELT), according to which proteins have evolved to fold along a smooth, funneled free energy landscape. Such “folding funnel” determines the efficient and reproducible collapse of the denatured polymer chain to its compact and functional 3D structure(29). The majority of GōMs employ a Coarse-Grained (CG) representation of the protein in which each residue is mapped onto a sphere centered at the position of the *C_α_* atoms. The residues in contact in the native state interact via attractive pair potentials, defined so that the energy minimum of the model corresponds to the native fold. This picture assumes that folding is dominated by native contact interactions while non-native interactions play a minor role(30). GōMs have been validated using both experimental data and more detailed simulation models(31–36), and their predictions are considered reliable in the framework of small protein folding.

GōMs have been widely used to study the folding of entangled proteins, providing valuable indications on their thermodynamics and kinetics (37–41), also in the framework of lasso folding(20, 22, 23). However, the underlying theory clashes with the presence of knots in proteins, as the formation of entanglements implies a high degree of coordination at different length scales which can hardly be encoded in native contact potentials. For example, the mandatory passage through the *specific, non-alternative* folding intermediates imposed by topological barriers, can trigger the untimely formation of native contacts, which can entrap the molecule in misfolded states. When this happens, the protein has to break such contacts and retrace the proper folding route. On the one hand this “backtracking” process can explain the longer folding times measured for knotted proteins, on the other hand it lowers the capability of GōMs to fold reproducibly, resulting in very low success rates(40).

For this reason the possibility of including non-native interactions within GōMs has been explored, obtaining significant improvements in the folding efficiency(42–45). This suggests that non-native interactions can play a crucial role in topologically complex folding, regulating the timing of native contacts formation, and guiding the concerted non-local moves required for the tying of the backbone(46). Moreover, in agreement with ELT, the folding of GōMs exhibits multiple pathways reaching the folded state(38, 42, 47), differently from the indications of all-atom MD(24) and experiments(18), which suggest the reproducible selection of a single route.

The presence of a dominant pathway can indicate that evolution has optimized knotted proteins in their folding behavior, minimizing the probability of misfolding, and promoting the most reliable and fast folding routes. Building on this optimality principle, in Ref. (48) an alternative CG description for the study of knotted folded proteins has been proposed. This model, dubbed Elastic Folder Model (EFM), is a CG, minimalistic description in which the folding of the polypeptide is driven exclusively by backbone bending and torsion potentials. EFM embodies the idea that the folding process has been kinetically optimized by evolution, in that it promotes the most efficient pathways of the backbone across the topological bottlenecks of knotted folding. To attain this optimality, once a specific protein is chosen, the relative magnitudes of its angular forces are tuned via a stochastic process, aimed at maximizing the folding success rate. The heterogeneous force-field obtained through this optimization procedure represents a sort of mean-field approximation of the cooperation between native and non-native interactions, and can provide valuable information on the folding mechanisms of the system under examination. This model has been used to investigate the folding of two small knotted proteins(48), observing a qualitative agreement with the all-atom simulations results of Ref.(24).

In the present work we have employed EFM simulations to study the folding of a glycoprotein, Granulocyte-macrophage colony-stimulating factor, that exhibits a CL native state. We have extended the original EFM introducing contact interactions between those cysteines that form a disulfide bridge in the native conformation. This allowed us to simulate the folding in oxidizing conditions, assessing the differences with respect to the process in a reducing environment. The angular potentials of this protein model have been optimized with a newly implemented evolutionary strategy, that could tune the model to fold reproducibly and rapidly, avoiding kinetic traps and efficiently surpassing the topological bottleneck associated to the formation of the lasso. The resulting dynamics has been compared with that of a well-established GōM(26) with the purpose of enlightening the most efficient folding pathways, in relation with the topological state of the protein. To this aim we have also introduced and employed two topological variables that, building on the minimal surface analysis(19) and the Gauss linking number(49) methods, allow to monitor the evolution of the CL topology along the MD trajectory.

As a result we could outline a detailed picture of the folding scenario, demonstrating that the same, kinetically optimal mechanism dominates in both reducing and oxidizing conditions. This folding route, characterized by the formation of the cysteine bridge after the lasso topology, is supported both by the GōM simulations at the fastest folding temperature and by the optimized EFM. These results show how the principle of kinetic optimality can determine the selection of a single folding mechanism among the possible ones, and qualify the considered protein as an interesting testing ground for all-atom simulations or experimental study.

## METHODS

In this work we have studied the folding of Granulocyte-macrophage colony-stimulating factor, a monomeric glycoprotein that acts as growth factor for white blood cells. We shall refer to the protein by using the PDB code of its crystal structure, 2GMF(50). 2GMF is an helical cytokine formed by 127 residues, of which 121 are resolved in the PDB structure, shown in Fig. 1. As highlighted in figure, 2GMF forms two cysteine bridges, which we name *b*_1_, connecting residues 88 and 121, and *b*_2_, connecting residues 54 and 96. 2GMF is classified as *L*_2_ lasso structure, where the covalent loop formed by *b*_1_ is threaded by a 12 residue hairpin from residue 43 to residue 53. Instead, *b*_2_ does not determine any lasso topology.

**Figure 1:**
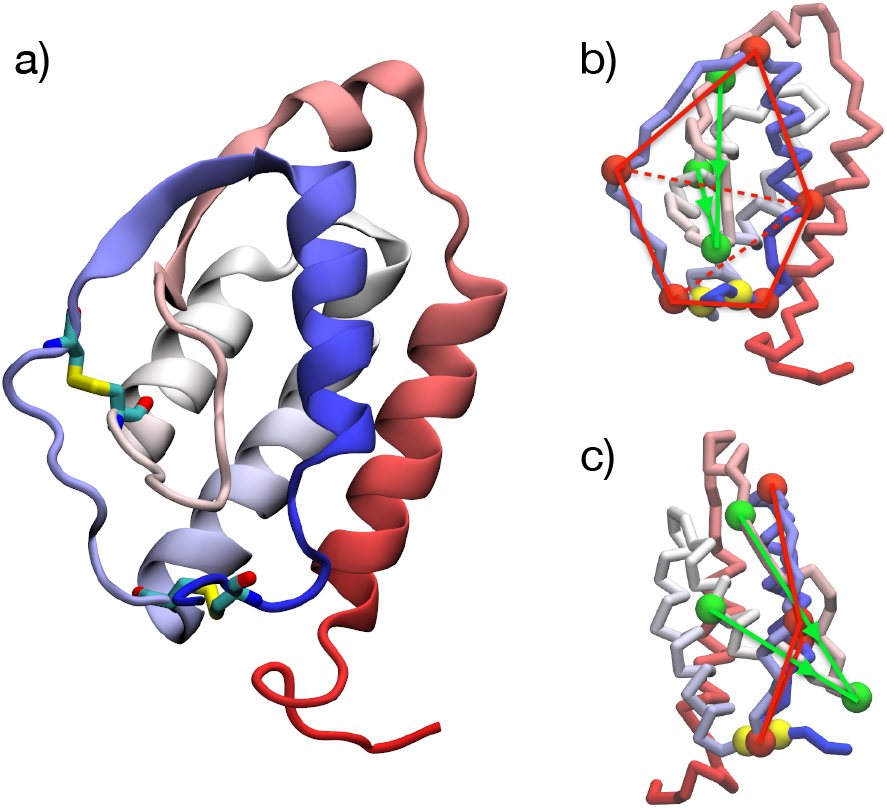
2GMF Protein structure and geometry of topological descriptors. Panel a): Cartoon representation of Chain A of 2GMF in its native fold. The cysteine bridges are shown with atomistic resolution. Panel b): view of 2GMF native structure, showing only the *C_α_* residues. Cysteines 88 and 121, forming *b*_1_, are represented as yellow beads. The structure reduction employed for the definition of the topological variables *L* and *G* (see the text) is also displayed: the 5 residues chosen to represent the loop are highlighted as red circles connected by red lines, while the 3 residues representing the threading hairpin, are highlighted as green circles connected by green lines. The red dashed lines indicate how the loop surface is divided in 3 triangles for computing *L*. The green arrows represent the integration verse along the hairpin segments used for the calculation of *G*. Panel c): same as b) but rotated. The coloring of backbone residues depends on their index along the chain, going from red (N-terminal) to blue (C-terminal).

Three different MD models of the protein were employed, a non-optimized EFM with homogeneous stiffness coefficients, an optimized EFM, obtained with the MFFO procedure presented in the following, and a GōM constructed using the native-contact based description proposed by Clementi et al. (26). The folding of 2GMF was studied by performing sets of MD runs starting from random stretched configurations, both in reducing and in oxidizing conditions. As discussed in the following, the stability of cysteine bridges in oxidizing environment is modeled in the EFM by means of native contact potentials between the cysteine pairs, and in the GōM by rescaling the existent native contacts. We shall employ natural units, indicating energies in units of ***ε***, temperatures in units of ***ε***/*k*_B_, lengths in units of ***σ***, and time-lengths in units of 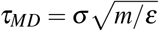, *m* being the bead mass.

### Elastic Folder Model Simulations

The EFM introduced in Ref. (48) is here reviewed in detail. The model describes an *N*-residues polypeptide chain by means of a CG representation, in which only the *C_α_* atom positions are retained, resulting in a chain of *N* identical beads connected by stiff bonds. The steric hindrance of each residue is represented by a short-range excluded volume interaction. As said, the driving force of the folding is modeled by bending and torsion potentials, parametrized so that the energy minimum is attained for a chosen reference configuration. In principle, this reference corresponds to the native PDB structure, however other choices can be convenient as well(48).

The total potential energy is:

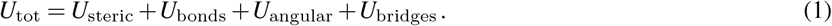

Weeks-Chandler-Anderson interaction(51) is used for the steric term:

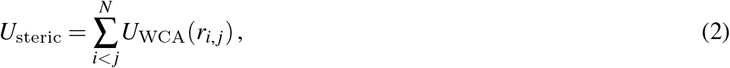

where *r_i,j_* = |**r**_*i*_ − **r**_*j*_| and the pair potential is given by:

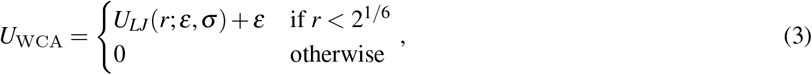

in which *U_LJ_* is the Lennard-Jones potential:

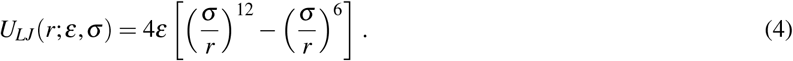

The chain beads are connected via Finitely Extensible Nonlinear Elastic (FENE) bonds(52), namely:

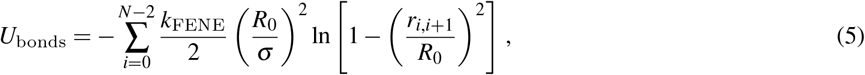

in which kFENE is the interaction strength parameter and *R*_0_ is the maximum bond length. The length scale ***σ*** is chosen equal to the steric diameter of Eq. (2), that corresponds to the separation between two consecutive *C_α_*, i.e. roughly 3.8 Å.

The remaining terms of Eq. (1) contain specific structural information of the protein which has to be described. As mentioned, the folding is guided by the angular potential, that generates the dynamics of the chain bending and torsion angles:

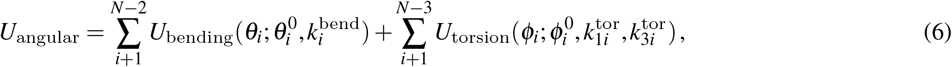

in which 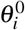 and 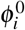 are, respectively, the *i*-th bending and torsion angles of the reference conformation. 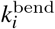 and 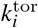 are the stiffness coefficients associated to the angular potentials, which are given by:

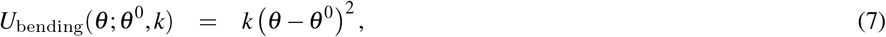

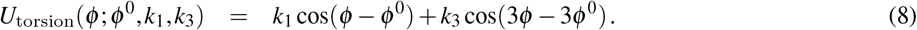

In EFM we consider a single torsion coefficient, imposing *k*_3_ = *k*_1_/3. Angular interactions such as Eqs. 7 and 8 (or analogous chiral potentials) have been included in Gō models as well(26, 27), in order to bias the formation of proper backbone chirality (53).

In this work we have modeled the formation of disulfide bridges by introducing an attractive potential term *U*_bridge_ between those *n*_B_ cysteine pairs {*c*_1_, *c*_2_} that form a bridge in the reference state.

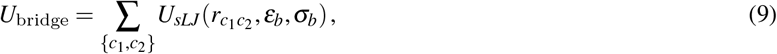

in which *r*_*c*_1_*c*_2__ is the distance between the cysteines and *U_sLJ_* is a truncated and force-shifted LJ potential, given by:

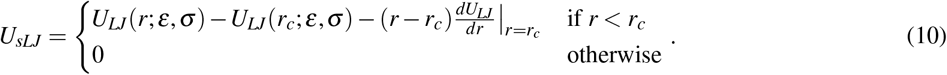

The scale length ***σ***_*b*_ is chosen so that the minimum of *U*_bridge_ is located at the reference distance between the residues in the considered pair.

The folding of this protein model is studied simulating its Langevin dynamics starting from a stretched (i.e. end-to-end distance ~ *N****σ***), randomly generated configuration. The potential parameters, as well as the MD settings, were chosen following the previous work on EFM(48). The FENE parameters had the typical values *k*_FENE_ = 30 and *R*_0_ = 1.5, the friction time of Langevin equation was ***τ***_frict_ = 1.0 and the integration time step was Δ*t* = 5 × 10^−4^. The EFM dynamics was integrated by means of an inhouse software.

### Single Force-Field Optimization

To satisfy the principle of optimality of the folding pathway, the EFM angular force parameters 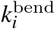 and 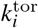 are tuned to maximize the success rate of the folding. In Ref. (48) this optimization is performed through a stochastic search procedure, which we recall here. Let us first define:

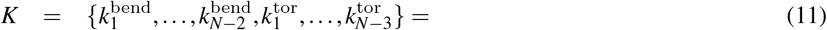

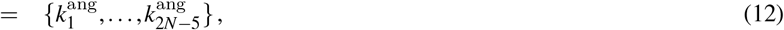

in which 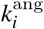 is used for both bending and torsion stiffnesses. We shall refer to *K* as the *force-field* of the model. The optimization step consists in two operations: first, a mutated force-field *K*′ is generated:

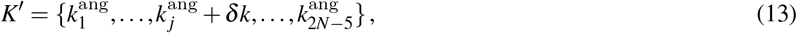

in which the *j*-th coefficient is modified by adding *δk. j* is randomly chosen among the 2*N* − 5 coefficients, while *δk* is generated with a prescribed probability distribution (e.g. in our calculation it is normally distributed, with standard deviation equal to 2.5). Second, the mutation to *K*′ is accepted or rejected according to a Metropolis-like criterion: *K*′ is tested by performing a set of *n* parallel folding simulations, starting from a randomly generated stretched configuration, and running for some properly chosen length ***τ***_run_. The outcome of the *n* test trajectories is then assessed by measuring 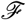, namely the Mean Square Displacement (MSD) from the target configuration **R**^0^, defined as:

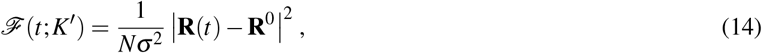

where **R**(*t*) is the configuration vector of the protein model, and |·| is the Euclidean distance. We then define:

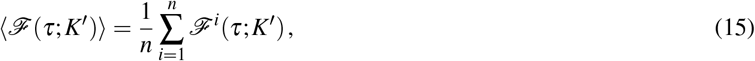

which is the average MSD computed at *t* = ***τ*** over the *n* test runs. ***τ*** is chosen so that Eq. 15 provides a measure of the folding success of the test runs. It can be set e.g. equal to ***τ***_run_ or, as in Ref.(48), chosen according to the the convergence of the MSD value along the trajectory. In the present work we have selected ***τ*** = ***τ***_min_, namely the time at which the MSD reaches its minimum value during the test run. The probability of acceptance of the new force-field *K*′ is then:

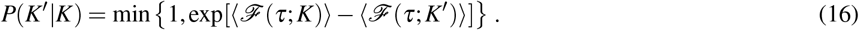

The operation just described is then iterated to minimize 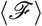, enhancing the average success rate of the folding trajectories. A schematic representation of this procedure, that we name Single Force Field Optimization (SFFO), is displayed in Fig. 2.

**Figure 2:**
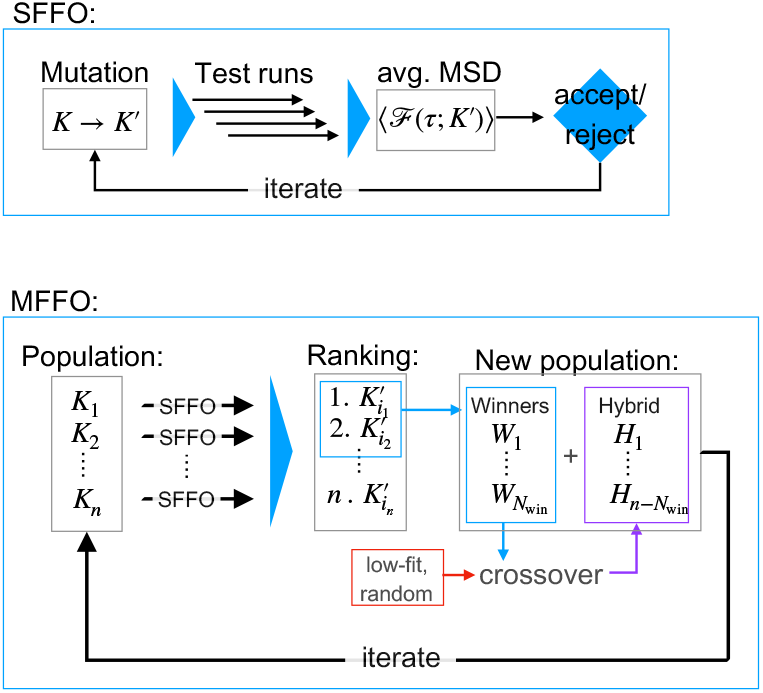
Schematic illustration of the SFFO and MFFO optimization algorithms.

For a polypeptide such as the smallest knotted protein MJ0366, with *N* = 82 residues, the parameter space is quite large and the SFFO algorithm can explore only a minimal portion of it in reasonable computation time. The situation can be partially improved by constraining the *k*^ang^ to be locally equal. For example, in Ref. (48) as well as in the present work, neighboring pairs of coefficients are constrained, that is:

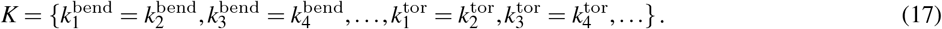

These local constraints reduce the dimensionality of the stochastic search, but also the generality of the model. In the present work we have employed Eq. 17, pairing neighboring angular coefficients.

### Multiple Force-Field Optimization

In this work we have employed a development of the SFFO strategy, aiming at a more efficient exploration of the *K*-space. The basic idea is to apply SFFO for the parallel optimization of several force-fields and then combine the results with an evolutionary strategy, as graphically illustrated in Fig. 2. An initial set, or population, of force-fields 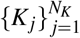 is chosen, and each of them undergoes *m* SFFO steps independently from the others. The resulting *N_K_* mutated force-fields are then ranked according to their capability of folding. The specific ranking criterion is discussed in detail later on. The *N*_win_ top-ranked force-fields, which we shall call “winners”, are selected to build the new population 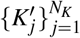, while the remaining, low-ranked candidates are discarded. The new force-field population is given by:

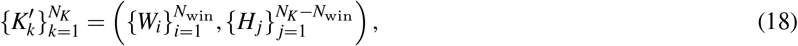

in which *W* indicates the winners, and *H* indicates a set of *N_K_* − *N*_win_ newly generated force-fields, which we shall refer to as “hybrid”. The latter ones are obtained by means of a *crossover* operation, typical of genetic algorithms (see e.g. Ref. (54)). More in detail, the *H_j_* are generated by combining fragments of force-fields, randomly picked from a set of parent force-fields, as displayed in Fig. 3). The parent set is formed by the *N*_win_ winners together with *N*_low_ “low-fit” candidates, that ensure diversity among the population. The latter can be selected among the worst ranked force-fields or, otherwise, generated with randomly distributed angular coefficients. Further details about the crossover operation are provided in the Supplementary Material (SM). Once the new population is set the optimization cycle is completed and the algorithm is re-iterated. We name this procedure Multiple Force Field Optimization (MFFO).

**Figure 3:**
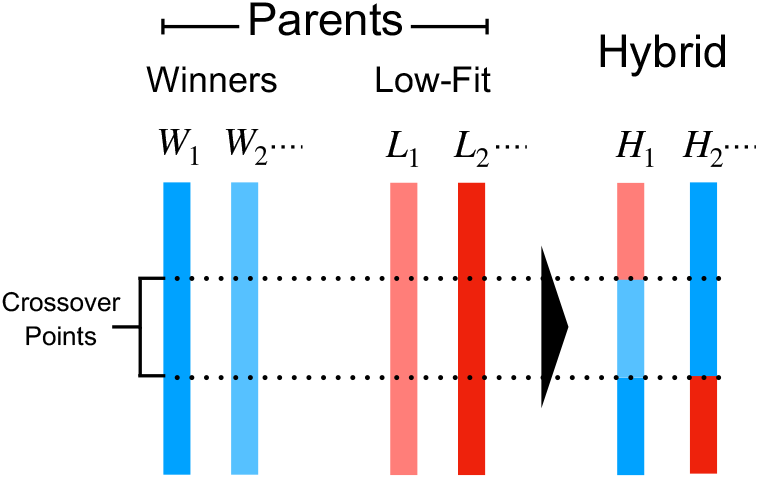
Schematic representation of the crossover operation generating the hybrid force-fields. The color bars indicate the sets of *k*^ang^ coefficients associated to the Winners and the Low fit force-fields. These are mixed randomly in the hybrid force-fields.

We now discuss the criterion for the force-field ranking, which naturally builds on the outcomes of the folding tests gathered during the SFFO steps. As explained, each SFFO mutation is tested via *n* folding simulations. The resulting *n* trajectories can provide indications on the folding propensities of the *N_K_* force-fields. One can e.g. compare the average MSD (Eq. 15) attained by each force-field. Another possibility, which we have adopted in the present work, is to rank the *N_K_* candidates according to *P_f_*, namely the folding probability along the test runs. More precisely, we have defined an estimate ***π***_*f*_ of the folding probability, based on the measurement of the MSD along the test runs. A threshold value 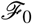 has been set, below which the protein is considered to be in the native state. Then, for a set of *n* test runs, we have defined:

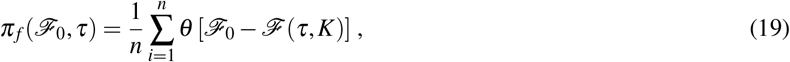

where ***θ*** is a function that switches from 0 to 1 when its argument becomes positive. In particular we used a Fermi function

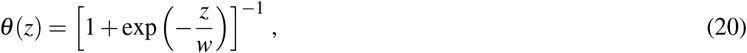

that switches continuously with length-scale w. Clearly, ***π***_*f*_ represents only a proxy of the real folding probability, on the one hand because the sole MSD is not always reliable in discriminating between the native basin and misfolded configurations, and on the other hand because it depends on a limited number of finite trajectories. As mentioned, the ranking operation has performed every *m* SFFO iterations. Therefore, the trajectories employed in computing Eq. (19) come from the *m*-th iteration. However, we can assume that the local mutations tested along each SFFO step have a relatively small effect on the force-field folding propensity. It is thus convenient to include in the ranking also the information from the previous *m* − 1 SFFO steps. To achieve this we have employed an exponential moving average, defined by the iterative formula:

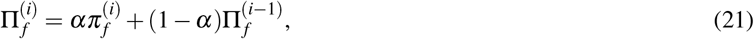

where 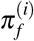 is the folding probability relative to the *i*-th SFFO iteration and *α* is the smoothing factor 0 < *α* < 1. The final value, i.e. 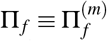, includes the contribution of all *m* SFFO iterations, assigning them a weight that increases exponentially with *i*. Thus the *N_K_* force-field candidates have been ranked by increasing values of Π*f*.

In the optimization presented in this work, the MFFO strategy has been applied to a population of *N_K_* = 16 force-fields, initially having homogeneous angular coefficients 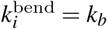 and 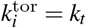, where *k_b_* and *k_t_* were chosen among the possible combinations of 20.0, 40.0, 60.0 or 80.0. Each force-field was optimized via SFFO, during which it mutated point-wise. The local mutations were accepted via a Metropolis criterion, based on the average MSD of 16 parallel folding trajectories (Eqs. 15 and 16) of length ***τ***_run_ = 3.5 × 10^3^. This trajectory time-length has been chosen based on the folding times measured for the HM, in order to promote only the faster folding routes. Every *m* = 50 steps the force-fields were ranked according to the value of Π_*f*_, as given by Eqs. 19 and 21, where the thresold MSD was 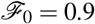, the switching length-scale *w* = 0.2 and the smoothing factor ***α*** = 0.03, corresponding to a decay time of ***τ***_*α*_ = 33 steps of the exponential moving average weight. As mentioned, the resulting Π_*f*_ is a proxy of the success probability *P_f_*, that provided an on-the-fly estimate of the optimization progress. After the force-fields were ranked, the 6 best were chosen as winners, and continued the optimization. The remaining 10 force-fields were constructed combining the winners and 4 randomly generated forcefields, with 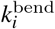 and 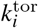 uniformly distributed between 30.0 and 60.0 (more details are reported in the SM).

### Gō Model Simulations

The employed Gō model is that introduced by Clementi et al. in Ref. (26). The system setup was generated using the SMOG web server (http://smog-server.org) (55). Details on the interaction potential, which is based on 12-10 Lennard Jones native contacts, can be found in the cited references. The shadow contact map(56) is used for the definition of native contacts. As mentioned before, also this description models the backbone stiffness with the angular potentials of Eq. 6. The stiffness coefficients are here homogeneous, set to *k*^bend^ = 40.0, 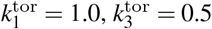. The formation of cysteine bridges in oxidizing condition is modeled by increasing the amplitude ***ε***_*ij*_ of the native contact potential associated to the cysteine-cysteine contacts. The value was set to ***ε***_*ij*_ = 10*k*_B_*T*, so that thermal fluctuations would hardly break the bridge once formed.

As for the EFM, the GōM folding is studied by means of Langevin dynamics, starting from random stretched configurations. GROMACS 2018.3 package(57, 58) was used for integrating the motion. The MD parameters were chosen consistently with the EFM simulations, with time step Δ*t* = 5 × 10^−4^ and friction time ***τ***_frict_ = 1.0. To select the simulation temperature we have performed a study of folding times and probabilities at different values of *T*, the results are presented in the SM.

### Topology Analysis

Minimalistic, CG models make it possible to collect a large statistics of folding trajectories, even in complex folding processes as those of self-entangled proteins. However, in order to gather useful information on the folding dynamics, the analysis of these trajectories strongly benefits from the definition of proper topological descriptors. Many methods for detecting the entangled state of a polymer chain have been proposed (see e.g. Refs. (59) for further details) and extensively applied. For example, in the framework of knotted proteins, knot searching algorithms have been used to classify the topology of known native structures, gathering a comprehensive database(3). In general these techniques operate on the three dimensional structure of a polymer chain, first by associating it to an equivalent closed curve(60, 61), so that the topological state is mathematically well-defined, and second by simplifying this structured curve without changing its topology(8, 62). The resulting curve is then analyzed by computing topological invariants(63, 64) and its entangled state is classified.

Although this is the typical approach used to analyze knotted proteins, the non-trivial topology recognized in CL structures, is not yet classified from a mathematical point of view(2). In Ref. (19) an approach specifically aimed at detecting CLs is presented. This technique, named Minimal Surface Analysis, uses triangulation algorithms mutuated from computer-graphics, to determine the minimal area surface spanned by a protein covalent loop. When this surface is obtained the lasso-type is detected by searching for segments of the backbone that pierce the minimal surface. This is a robust method to assess and classify CL structures, and it has been employed to establish a database of polymeric structures characterized by this topology(19, 21). However in the present work we are interested in descriptors that can monitor the topological state along the folding trajectory of a specific protein. To this purpose the computation can be expensive, and a faster, less general method could be more effective.

We can thus exploit the fact that proteins fold reproducibly in a well-defined topology, which is known a priori. For this reason we relax the generality of the topological descriptor, and focus on the specific native geometry of the system under consideration. In CL geometries the main topological feature is a covalent loop closed by a cysteine bridge, pierced by part of the backbone. For simplicity, we limit the discussion to the case of a single loop and a single threading segment, the strategy can be then generalized to more complex topologies. Let *l*_1_,…, *l_N_l__* be the indexes of loop residues and *t*_1_,…, *t_N_t__* be the indexes of the threading segment residues. We operate a *reduction* of the structure, selecting only few crucial residues, namely 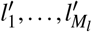 for the loop and 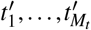 for the threading tail, where *M_l_* < *N_l_* and *M_t_* < *N_t_*. The residues *l*′ and *t*′ are chosen so that their position can describe whether the protein is in the native topology. This operation is similar to the smoothing performed for protein knot detection(65), however the procedure is not automated, and needs some preliminary analysis of the structure and folding behavior. For clarity, in Fig. 1 (and in the SM) the reduction we adopted for 2GMF is illustrated. The surface spanned by the *M_l_* loop residues is then approximated by *M_l_* − 2 triangles, with vertexes corresponding to the *l*′ residues positions. After this the threading of a |**R**_*t*′+1_ − **R**_*t*′_| segment through the loop can be verified by computing its intersections with the surface triangles. Once the number and directions of the piercings through the loop surface are determined it is clear whether the protein has attained its native topology. By means of continuous switching functions (see e.g. Eq. 20) we can associate this binary information to a continuous value *L* varying from 0 (non-native topology) to 1 (native topology), we name this quantity *lasso-variable*. The approximated surface formed by the *M_l_* − 2 triangles is not the minimal area surface of Ref. (19), which is typically formed by many more triangles. However, in our study, this simplification is convenient to speed up the calculations.

Another interesting approach to topology detection is adopted in Refs. (49, 66). The idea developed in these works is that of employing the Gauss linking number(67), namely the double line integral:

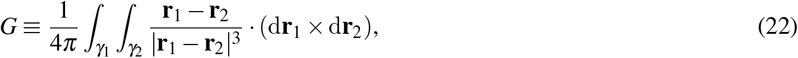

in which the integrals are performed along the two curves *γ*_1_ and *γ*_2_, **r**_1_ and **r**_2_ being the position vectors belonging to *γ*_1_ and *γ*_2_ respectively. If the two curves are closed (in ℝ^3^), *G* takes an integer value, that is a topological invariant typically used to define links. By applying a proper closure procedure, *G* can be therefore employed to detect the entanglement of two chains.

A crucial observation is that, when *γ*_1_ and *γ*_2_ are not closed, *G* is not an integer topological invariant, but it still provides relevant information on the curves’ mutual entanglement(67). This property can been then exploited to assess the linking in protein dimers(49), or the self-entanglement of folded proteins(66) without the need to define a closure operation. A strong correlation has been found between the value of *G* computed over open curves and its “closed counterpart”. This indicates that Eq. 22 can be used as a descriptor for the topological state of entangled structures such as CLs. Once again, since we are interested only in a specific topological state, we have simplified the calculation of *G* in the same way as done for *L*. Therefore we have computed *G* by applying Eq. 22 to the polygonal curves defined by the *M_t_* and *M_l_* residues selected by structure reduction. In this case however, we have adopted the convention that the bridge-forming cysteines are always the ends of the integration along the covalent loop. This way, *G* depends on the distance between the two cysteines, being affected to the opening and closing of the covalent loop.

The cross product in Eq. 22 implies that *G* depend on the relative orientation of *γ*_1_ and *γ*_2_ curves. Therefore one has to define an orientation along which the two sub-chains are integrated. In the present work we have not fixed any conventional orientation, since we have not compared different molecules. However, we have computed *G* for an *L*_2_ lasso structure, in which the tail pierces the loop twice, in opposite directions (as shown in Fig. 1). In this case, the contribution to *G* provided by the threading in one direction is partially compensated (or entirely compensated, if the curves are closed) by the threading in the opposite direction. To adapt *G* such that it can detect this double piercing we have separated the threading tail in two parts, assigning two different orientations for the calculation of Eq. 22. As a result, the contributions coming from the two piercings add up, detecting the *L*_2_ state.

## RESULTS

### Homogeneous EFM

We first report the folding behavior of 2GMF described by an homogeneous EFM, in which the angular potentials (see Eq. 1 in Sec. Methods) are parametrized using homogeneous angular coefficients 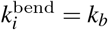 and 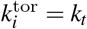, where *k_b_* = 36.5 and *k_t_* = 38.5. From now on we shall refer to this representation as Homogeneous Model (HM). The order of magnitude of *k_b_* and *k_t_* is consistent with the settings used in Ref. (48), but the values were chosen equal to the average of the optimized bending and torsion coefficients, presented in the following. This choice allowed us to assess the impact of the force-field heterogeneity introduced by the optimization procedure. Consistently with Ref. (48), we have studied the model at *T* = 0.1, that is below the melting point of the model and, as shown in the following, determines a quite frustrated free-energy landscape. An ensemble of 2048 folding trajectories has been collected, both in reducing and oxidizing conditions. Eq. 9 was used to model the bridge in oxidizing environment.

In order to define the folding criterion we monitored two variables, the Root Mean Square Displacement (RMSD) 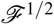 from the native state, and the lasso variable *L*, indicating the formation of the CL topology (see the Methods section for the definitions of 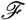 and *L*). We have selected two threshold values for 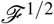 and *L*, considering the protein as fully folded only if both 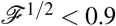 and *L* > 0.9. In most of the cases the RMSD criterion was sufficient to classify the nativeness, however the measurement of *L* has allowed to point out few false positives, and to distinguish successful folding trajectories with better accuracy. Once the success criterion has been defined, the probability of folding was estimated as *P_f_* = *n*_f_/*n*_tot_ where *n*_f_ is the number of trajectories attaining the folding, and *n*_tot_ = 2048 is the total number of runs. This estimate of the success rate depends on the length τ_run_ of the simulated trajectories. Since the EFM focuses on the optimal pathways of folding we aimed at observing those folding events that occur within the initial stages of the dynamics, not long after the collapse of the polymer chain. We have chosen τ_run_ = 1.5 × 10^4^ which, as shown in the following, is enough to capture all the fastest folding events obtaining indications on the timescales of the slower processes as well.

The computed *P_f_* of HM in reducing conditions is equal to 55%, while in oxidizing conditions the folded configuration is reached by the 17% of the trajectories. This shows that the topological barrier introduced by the cysteine bridge significantly increases the frustration of the model. We define the “folding landscape” as *F* = −log *f*, where *f* is the frequency histogram of some chosen reaction variables (e.g. the RMSD), computed over the ensemble of trajectories. This quantity is sometimes named “non-equilibrium free-energy surface”(68,69). We also introduce the “successful folding landscape” *F_s_* = −log *f_s_*, which considers only those trajectories that reach the native state.

In Figs. 4A and B the folding landscape of the HM in reducing and oxidizing conditions is reported, as a function of the RMSD and of *d*_*b*_1__, the distance of the two cysteine residues forming *b*_1_. The corresponding *F_s_* is instead shown in Figs. 4C and D. By comparing the successful trajectories to the whole ensemble we observed that the native basin is located in the region 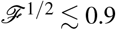. In both environmental conditions, the landscapes show a variety of metastable states, testifying the roughness of the free-energy surface. Since, except from the bridge potential, EFM does not introduce native contacts, this roughness is the result of the topological bottlenecks encountered during the folding trajectories. In particular, if we consider only the successful trajectories in reducing conditions (Fig. 4C), we observe a metastable state at RMSD~ 2.0, presumably connected to the native basin by an open-bridge pathwhay, with *d*_*b*1_ ~ 4.0. In oxidizing conditions (Fig. 4D) this metastable state is perturbed by the action of the bridge potential, which restrains part of the trajectories close to its minimum, where the covalent loop is closed. Most of these trajectories remain trapped in this state, and cannot overcome the topological barrier to reach the native basin.

**Figure 4:**
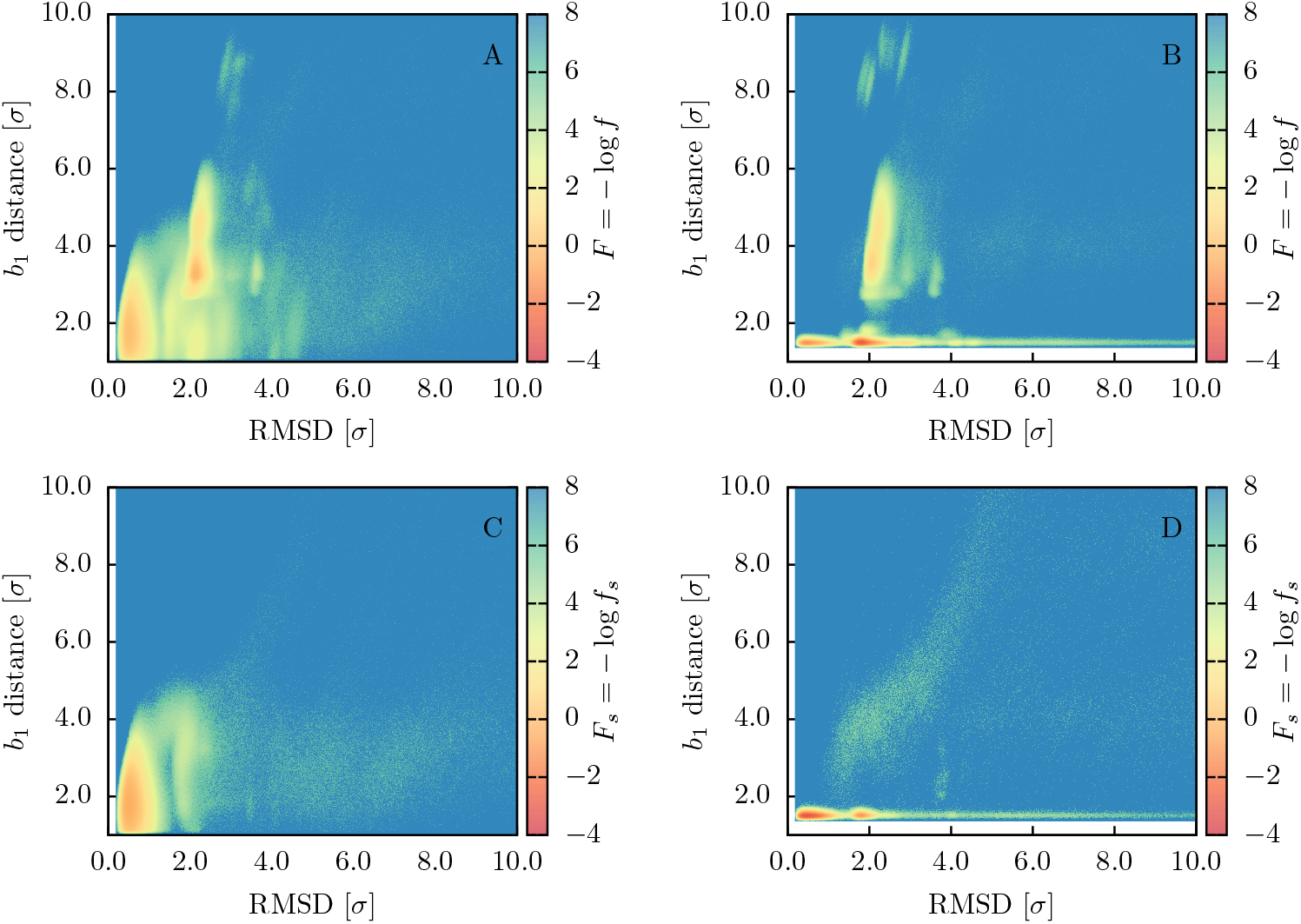
Folding landscapes *F* and *F_s_* of the HM as a function of the RMSD from the native structure and of the *b*_1_ bridge distance. Panel A: *N* = 2048 trajectories in reducing conditions. Panel B: *N* = 2048 trajectories in oxidizing conditions. Panel C: successful trajectories (*N* = 1133) in reducing conditions. Panel D: successful trajectories (*N* = 350) in oxidizing conditions.

In order to extract valuable information about the folding pathways, we have employed the lasso variable *L* and the Gauss’ linking number *G*, defined in the Methods Section. Both *L* and *G* are useful to monitor the topological state of the protein along the trajectory but, since they exhibit a different behavior, we employ them for different purposes. Since *L* switches sharply from 0 to 1 when the native topology is attained, it is used to detect the time of formation of the lasso and, as mentioned before, to assess the folded state. *G* displays instead a smoother behavior, it is thus employed as reaction variable for computing the folding landscape, as shown in Fig. 5, where *F_s_* (*G*, *d*_*b*1_) is reported. The plot confirms that in reducing conditions the model establishes the lasso topology (attaining *G* ≳ 1) while the loop is open, and that the metastable state preceding the folding can be identified with a populated region without lasso conformation (G ~ 0). In oxidizing conditions the topological barrier is instead surpassed along two separate pathways, either with closed or open loop. We can classify the folding pathways as follows:

1. A “threading” mechanism, in which the contact between C88 and C121 is formed before the topology, then the closed loop is threaded by the hairpin to reach the native basin.
2. A “bridge reopening” mechanism, in which, again, the covalent loop is closed before the lasso is formed. The topology is then attained in a second moment thanks to a wide fluctuation of the bridge distance, and to the subsequent penetration of the loop by the hairpin.
3. An “open loop” path, in which the lasso is formed before the contact between C88 and C121, with the loop that “wraps around” the hairpin to form the native state.

**Figure 5:**
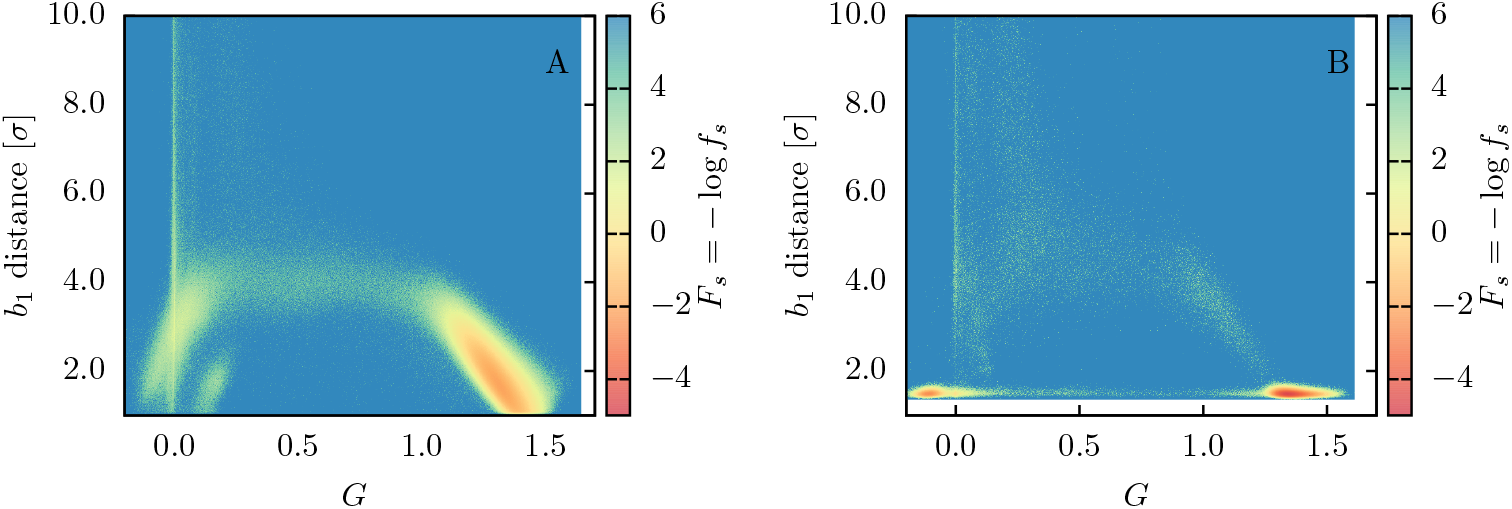
Successful Folding landscape *F_s_* of the HM as a function of the Gaussian linking number *G* and of the *b*_1_ bridge distance. Panel A: successful trajectories (*N* = 1133) in reducing conditions. Panel B: successful trajectories (*N* = 350) in oxidizing conditions.

A graphical illustration of these three processes is provided in Fig.6. The successful trajectories can be classified according to these three pathways, by performing a “kinematic” analysis that compares the timing of the main events in the folding process. For each trajectory we thus computed three transition times: a) the bridge formation time *t*_b_, namely the first time at which C88 and C121 approach at a distance *d*_*b*_1__ < 1.5*σ*_*b*_1__ = 1.992, b) the time of first topology formation *t_k_*, when *L* > 0.9, and c) the folding time *t_f_*, that is when the protein first visits the native basin (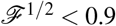 and *L* > 0.9). We required that the conditions for a), b) and c) remain valid for Δ*t* = 10 for the transition to be completed. Then, by comparing the measured *t_b_*, *t_k_* and *t_f_*with the time evolution of *d*_*b*_1__, which signals the closure of the loop, and of *L*, which indicates the topological state, we could classify the folding routes traveled by the protein in successful simulations.

**Figure 6:**
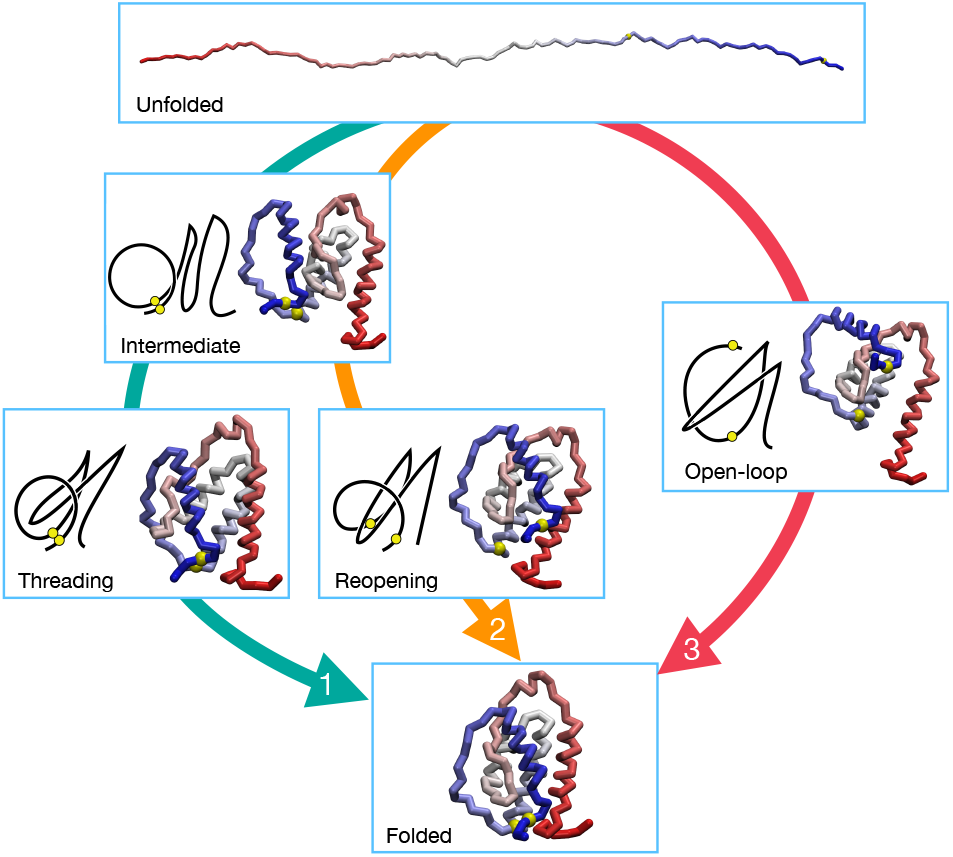
Illustration of the three folding pathways revealed by 2GMF EFM simulations. Each box contains the snapshot of a representative configuration along the corresponding folding route (represented by a colored arrow, numbered according to the pathway definition in the text). For further clarity, intermediate configurations are provided with a schematic diagram of the structure.

In Fig.7 the bridge formation times *t_b_* are plotted versus the folding times *t_f_* for each successful HM trajectory. The mechanism associated to each trajectory is indicated by different colors. The fraction of trajectories undertaking different routes is reported in Tab. 1. The folding mechanisms are differently distributed in reducing and oxidizing simulations. In the first case the successful folding events are similarly divided between open-loop and reopening pathway, while a relatively small number of threading trajectories is detected. Instead, in oxidizing conditions the reopening is prevented by the action of the cysteine bridge potential and, while the model mostly relies on the open-loop route, threading events are significant.

**Figure 7:**
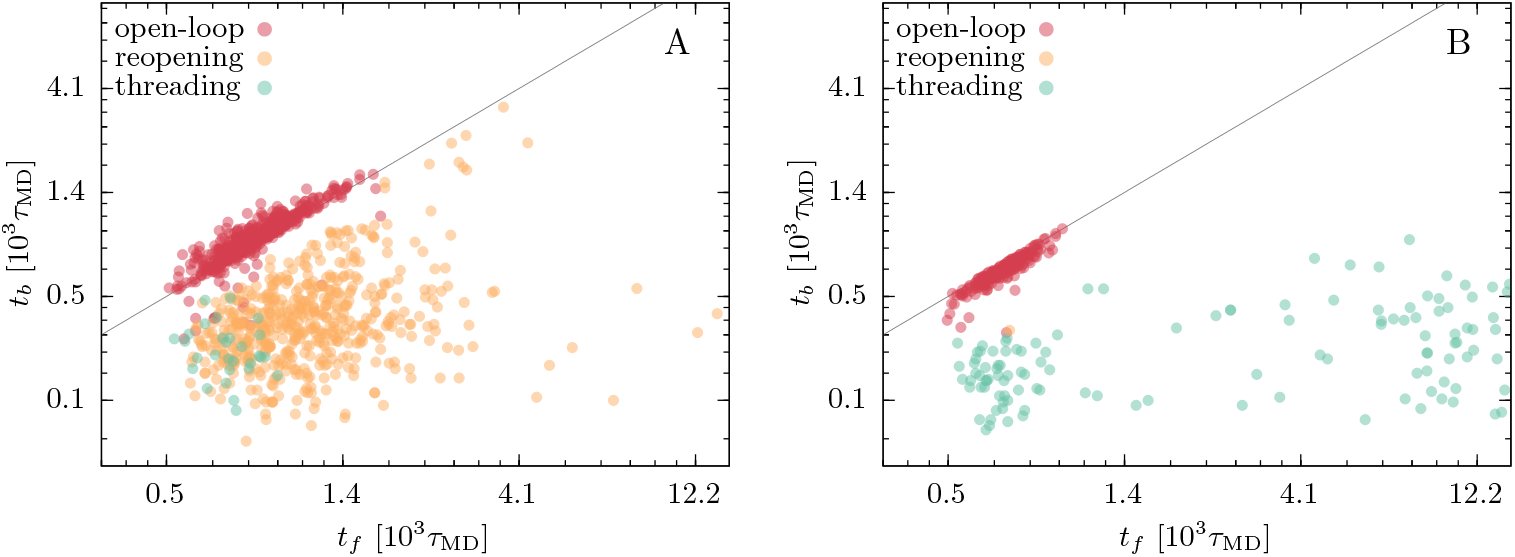
*t_b_* versus *t_f_* for the successful trajectories of the HM. Panel A shows the results of the *N* = 1133 successful trajectories in reducing conditions and panel B shows the results of the *N* = 350 successful trajectories in oxidizing conditions. The color of the circles indicates the folding pathway, following the classification indicated in the text. The black line corresponds to *t_b_* = *t_f_*.

**Table 1:**
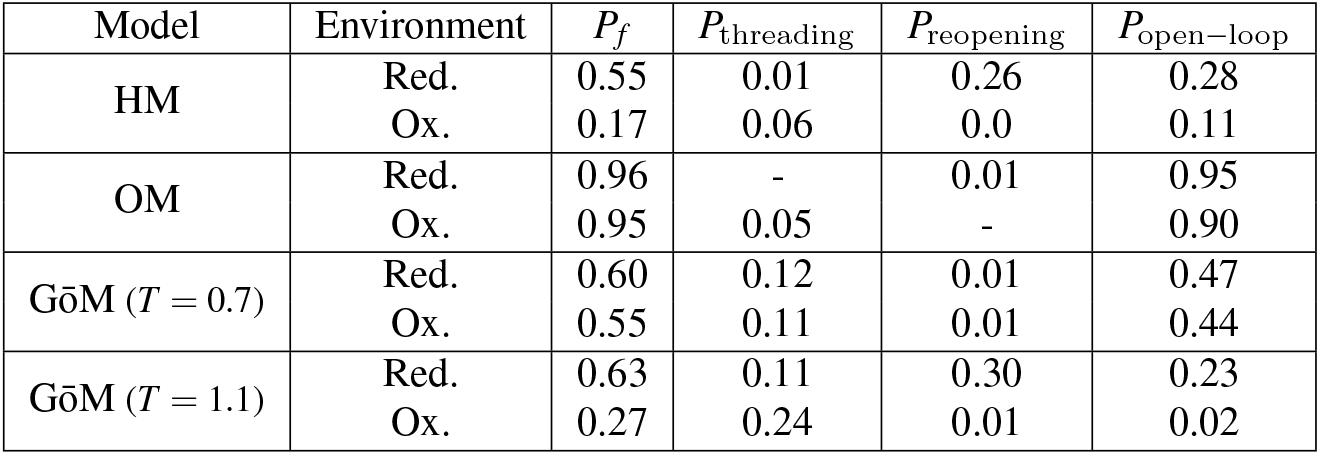
Probability of folding *P_f_* and pathway distribution. Folding probability *P_f_* and probability of undergoing different mechanisms (*P*_threading_, *P*_reopening_ and *P*_open–loop_), for each of the considered models, in reducing and oxidizing conditions. The probabilities are estimated as frequency of occurrence over 2048 trajectories of length ***τ***_run_= 1.5 × 10^4^.

| Model | Environment | $P_f$ | $P_{\text{threading}}$ | $P_{\text{reopening}}$ | $P_{\text{open-loop}}$ |
| --- | --- | --- | --- | --- | --- |
| HM | Red. | 0.55 | 0.01 | 0.26 | 0.28 |
|  | Ox. | 0.17 | 0.06 | 0.0 | 0.11 |
| OM | Red. | 0.96 | - | 0.01 | 0.95 |
|  | Ox. | 0.95 | 0.05 | - | 0.90 |
| GōM ($T = 0.7$) | Red. | 0.60 | 0.12 | 0.01 | 0.47 |
|  | Ox. | 0.55 | 0.11 | 0.01 | 0.44 |
| GōM ($T = 1.1$) | Red. | 0.63 | 0.11 | 0.30 | 0.23 |
|  | Ox. | 0.27 | 0.24 | 0.01 | 0.02 |

Another aspect that emerges from Fig. 7 concerns the folding time-scales characterizing the different pathways. Most of the observed folding events occurred for *t* < 10^3^, in particular those undergoing open-loop mechanism. In reducing conditions the re-opening events are distributed also beyond this time-scale, while the few threading events were faster. This is somehow counter-intuitive, as we expect that the entropic barrier of piercing the loop is larger when this is closed. If we look at the oxidized model results, we observe that threading events exhibit a bimodal time distribution, this suggests the existence of two possible threading pathways, a fast process, taking place for *t* < 10^3^ and a slower one, that requires a timescale comparable to τ_run_ = 1.5 × 10^4^. This bimodality disappears in reduced folding, in which the slow threadings are suppressed as the reopening of the bridge occurs over faster timescales.

Overall, this analysis reveals the main features of the folding of 2GMF as described by the EFM, and highlights the role of the topological barrier in selecting the accessible mechanisms to attain the native state. We underline here the importance of the defined topological diagnostics, *L* and *G*, in clarifying the folding pathway scenario.

### Optimized EFM

In this section we report the folding behavior of the EFM when optimized with MFFO, the evolutionary algorithm described in the Methods Section. As most of the lasso structures, 2GMF is a secreted protein, and its folding occurs in the endoplasmic reticulum, that is an oxidizing environment. For this reason the MFFO has been performed in oxidizing conditions.

The progress of the optimization procedure is displayed in Fig.8, where the evolution of the folding success rate is reported. We observe that, as the MFFO introduces heterogeneity in the angular interactions, the rate increases significantly reaching a value larger than 0.95. In Fig.8 we also show how the folding success rate evolved when no crossover between different force-fields was operated. This represents the success rate resulting from 16 independent SFFO runs (namely the serial stochastic optimization algorithm of Ref. (48)). It is evident that the MFFO approach provides a remarkable boost to the optimization, attaining a strong folding reproducibility, before the independent SFFOs exhibit any significant improvement. This substantial advancement in the optimization strategy opens the possibility of employing the EFM for the study of larger proteins, undergoing even more complex folding processes.

**Figure 8:**
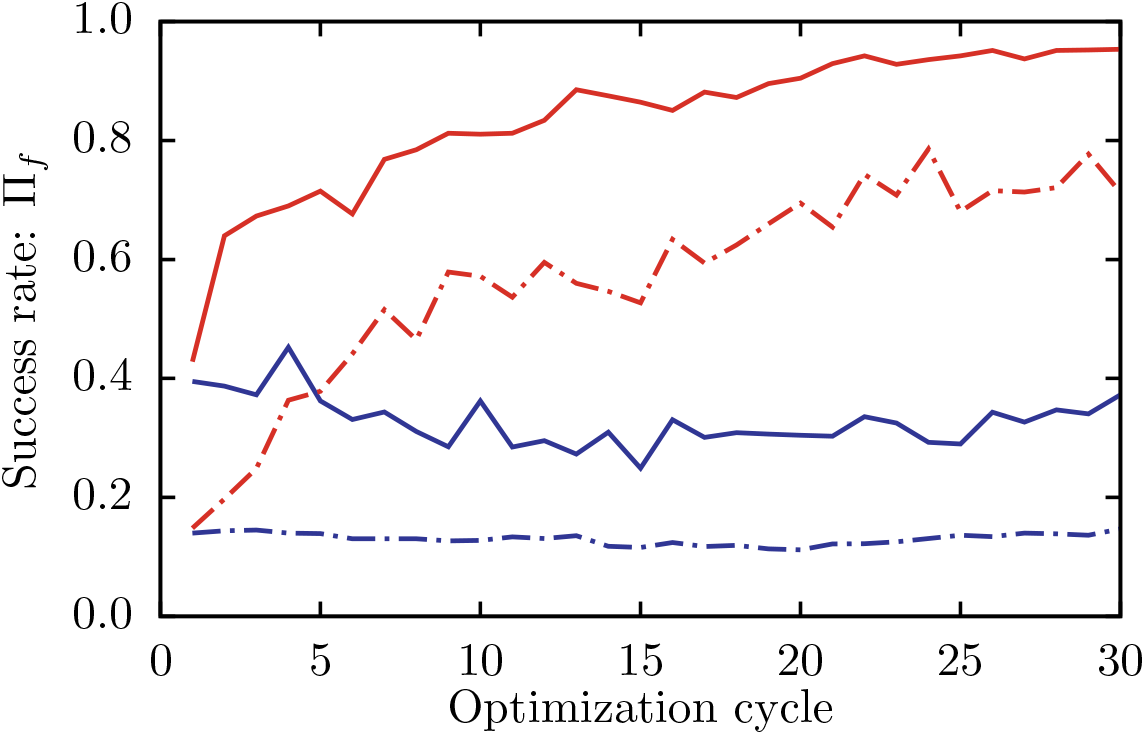
Average success rates of the folding trajectories performed during the optimization procedure. The reported quantity, Π_*f*_, is a proxy of the folding probability, the definition of which is provided in the Methods section. The red curves correspond to MFFO combining *N_K_* = 16 force-fields, the blue curves correspond to an MFFO without crossover of force-fields, equivalent to *N_K_* parallel SFFOs. Solid lines indicate the success rate of the best ranked force-field, while dot-dashed line indicates the average rate of the *N_K_* concurrent force-fields.

After 30 MFFO cycles we chose the top ranked force-field and tested it over 2048 folding trajectories, both in reducing and oxidizing conditions. We shall refer to this optimized model with the acronym OM. As described before, the bending and torsion stiffnesses of the HM have been set equal to the average values of the OM, this way we could assess the impact of heterogeneity on the folding behavior.

The *P_f_*’s obtained for the OM are reported in Tab. 1. We notice how the OM reaches high probabilities both in reducing and oxidized folding, showing that the heterogeneity of angular forces can be crucial to achieve a nontrivial topology in a reproducible way, in agreement with what found for knotted protein folding in Ref. (69). We then investigated the successful folding landscape *F_s_* associated to the OM, reported in Fig.9 as a function of 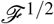 and *d*_*b*_1__, and in Fig. 10 as a function of *G*and *d*_*b*_1__. The landscapes look qualitatively different to those of Figs. 4 and 5, indicating that the OM selects different folding pathways with respect to HM. In particular we can appreciate how the non-entangled intermediate state is now less populated and how the closure of the cysteine bridge mostly occurs as a late event.

**Figure 9:**
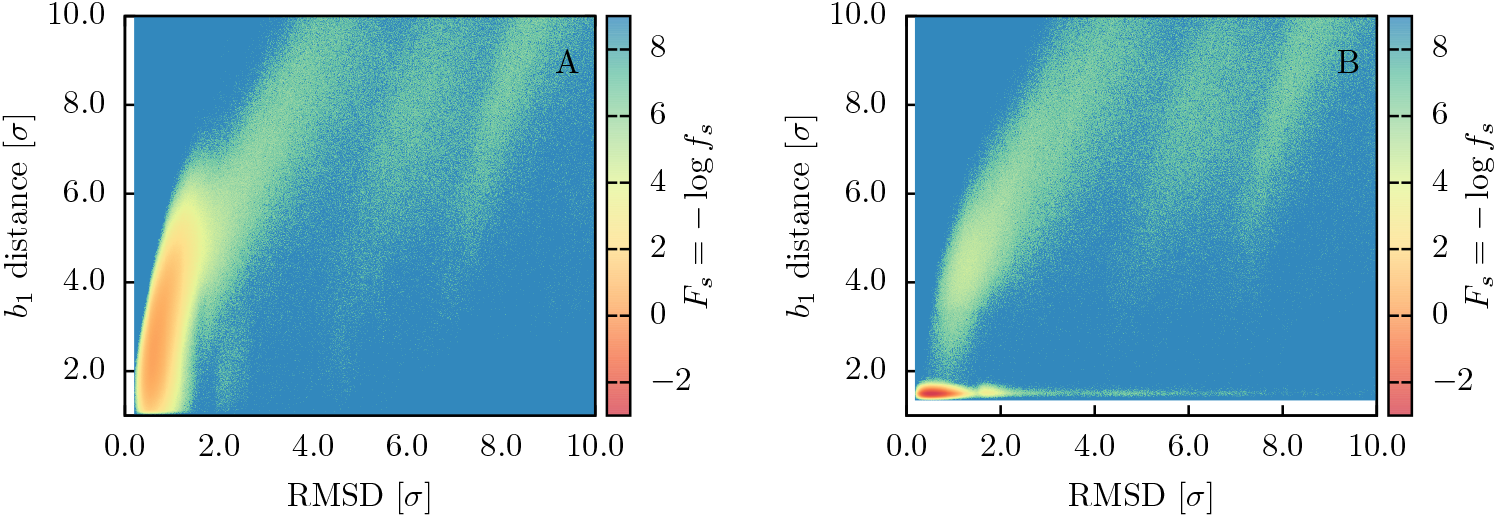
Successful Folding landscape *F_s_* of the OM as a function of the RMSD from the native structure and of the *b*_1_ bridge distance. Panel A: successful trajectories (*N* = 1970) in reducing conditions. Panel B: successful trajectories (*N* = 1946) in oxidizing conditions.

**Figure 10:**
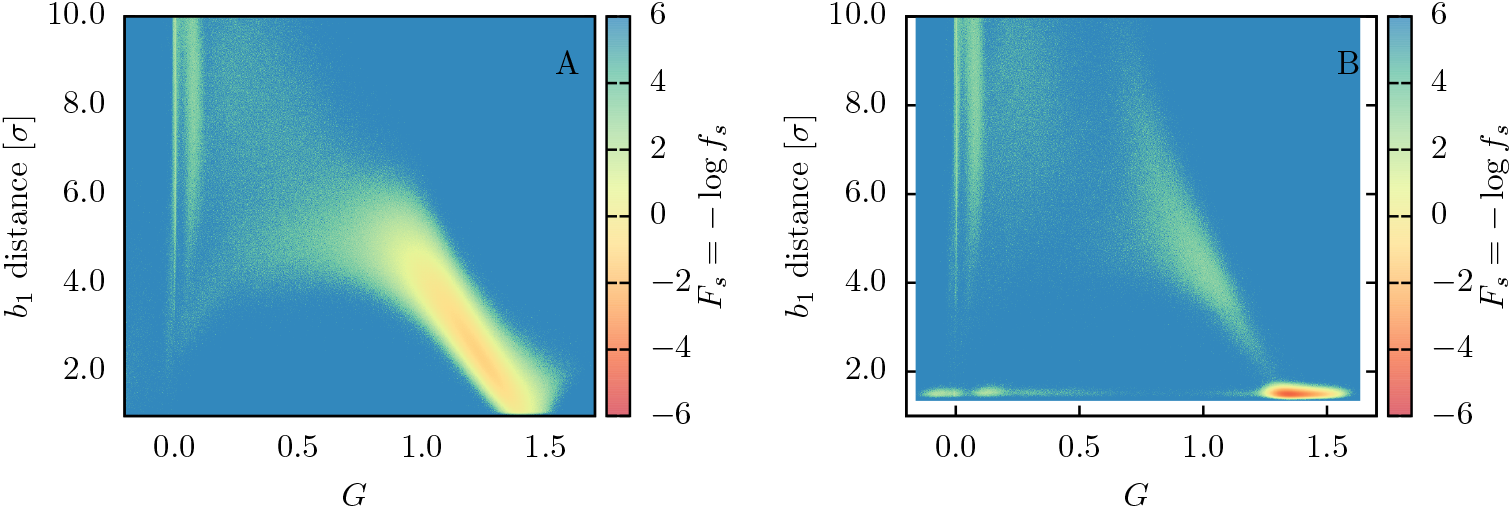
Successful Folding landscape *F_s_* of the HM as a function of the Gauss linking number *G* and of the *b*_1_ bridge distance. Panel A: successful trajectories (*N* = 1970) in reducing conditions. Panel B: successful trajectories (*N* = 1946) in oxidizing conditions.

To assess which folding pathways are more populated we repeated the kinematic analysis operated for the previous model. The results, shown in Fig. 11, reveal that the bridge formation and folding times are on average slower than in the HM model. The optimization acted on the timescale of the folding events by delaying the closure of the loop. As a result, the open-loop folding mechanism is promoted and characterizes the great majority of the trajectories, as indicated in Tab. 1. In EFM, the open-loop folding turns out to be the optimal route to the formation of the native lasso fold, in agreement with the intuition that the closure of the covalent loop determines an entropic barrier, slowing down the process. The behavior of OM shows how the optimization pressure, building on the requirement of a reproducible and efficient folding, can select a pathway among the possible ones, and polarize the mechanism of folding, similarly to what is observed in experiments and simulations of small, knotted protein folding(18, 24).

**Figure 11:**
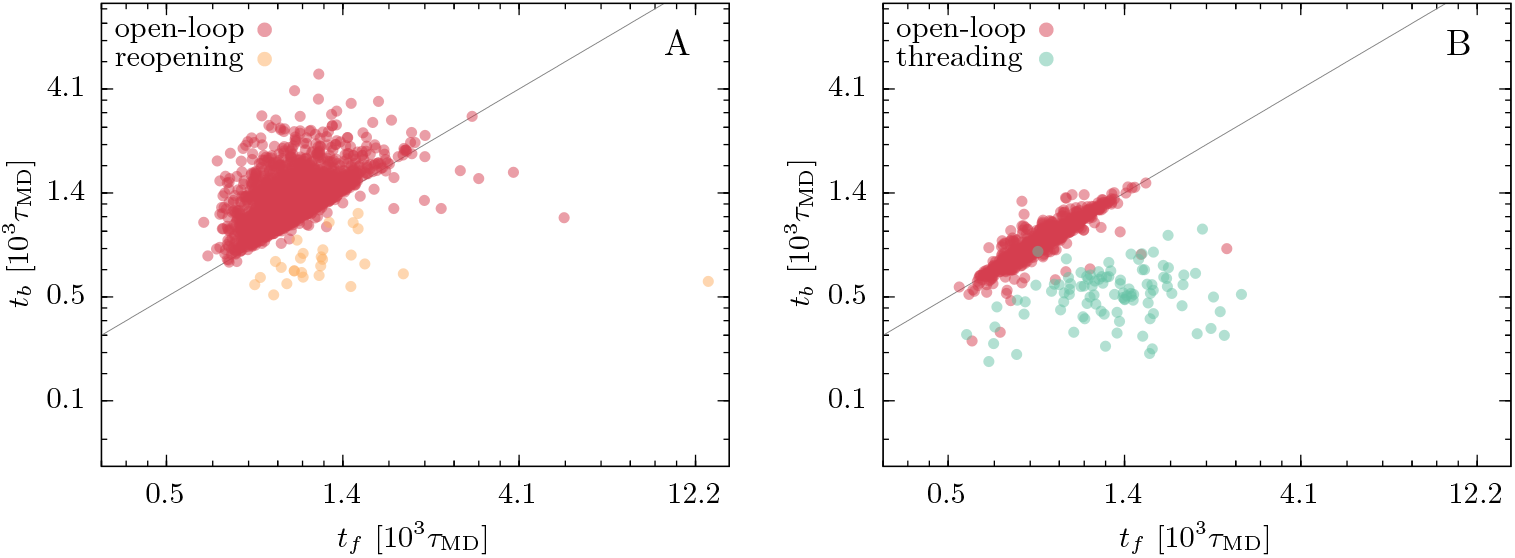
*t_b_* versus *t_f_* for the successful trajectories of the OM. Panel A shows the results of the *N* = 1969 successful trajectories in reducing conditions and panel B shows the results of the *N* = 1946 successful trajectories in oxidizing conditions. The color of the circles indicates the folding pathway, following the classification indicated in the text. The black line corresponds to *t_b_* = *t_f_*.

### Gō Model

To complement the picture obtained by means of the EFM we have performed a set of folding simulations employing the well-established GōM proposed by Clementi et al. (26), that has already been used by Haglund et al. to study lasso proteins(20, 22, 23). For details on the description we refer to the cited references, here we just underline that the native contacts establish through a 12-10 Lennard Jones potential, which is the main driving force of the folding. As mentioned in Sec. Methods, also this description models the backbone stiffness with the angular potentials of Eq. 6. The stiffness coefficients are homogeneous, set to *k*^bend^ = 40.0, 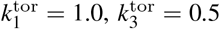. Following Ref. (70) we model the oxidizing conditions by rescaling the contact potential between the cysteines that form bridges in the native conformation. As a result of the temperature study presented in the SM, we have chosen to simulate this model at a temperature *T* = 0.7, at which the folding is kinetically optimal, or minimally frustrated(27, 35).

The folding criterion adopted for this model is the one chosen for EFM, namely requiring that 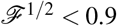 and *L* > 0.9 simultaneously. However, since the dihedral angles are substantially less stiff than in the EFM, the computation of *L* must involve a larger number of residues (see the SM for further details). The folding success rates resulting from 2048 simulations in both reducing and oxidizing conditions are reported in Tab. 1. The measured probability is in both cases above 0.55, with a lower value in oxidized conditions. This similarity in folding propensity suggests that the topological barrier imposed by the formation of the bridge does not have a substantial effect in this model. This is possibly related to the fact that the native contact between the cysteines is present also in the model under reducing conditions, albeit weaker. However, the analysis of the folding pathways provides further indications to explain this similar capability of folding.

Applying the same criteria employed for the EFM, we have analyzed the successful trajectories collected with reduced and oxidized GōMs, and assessed the population of different folding mechanisms. The results are reported in Tab. 1, and represented in the *t_b_* versus *t_f_* plots of Figs. 12A and B. The data indicate that the distribution of folding mechanisms is similar in reducing and oxidizing conditions. This symmetry confirms indeed that the successful folding events are not significantly affected by the cysteine bridge potential. However, most of the trajectories adopted an open-loop pathway, in which the topology forms before the contact of cysteine residues. This mechanism selection is the main reason why the model folds with a similar success rate in both environmental conditions. Moreover, we have found that the behavior of GōM, at the temperature of fastest folding, is in qualitative agreement with that of OM. Indeed both descriptions select a folding pathway characterized by the formation of the CL topology before the closure of the covalent loop, pointing at this way of overcoming the topological bottleneck as the most efficient option for the protein.

**Figure 12:**
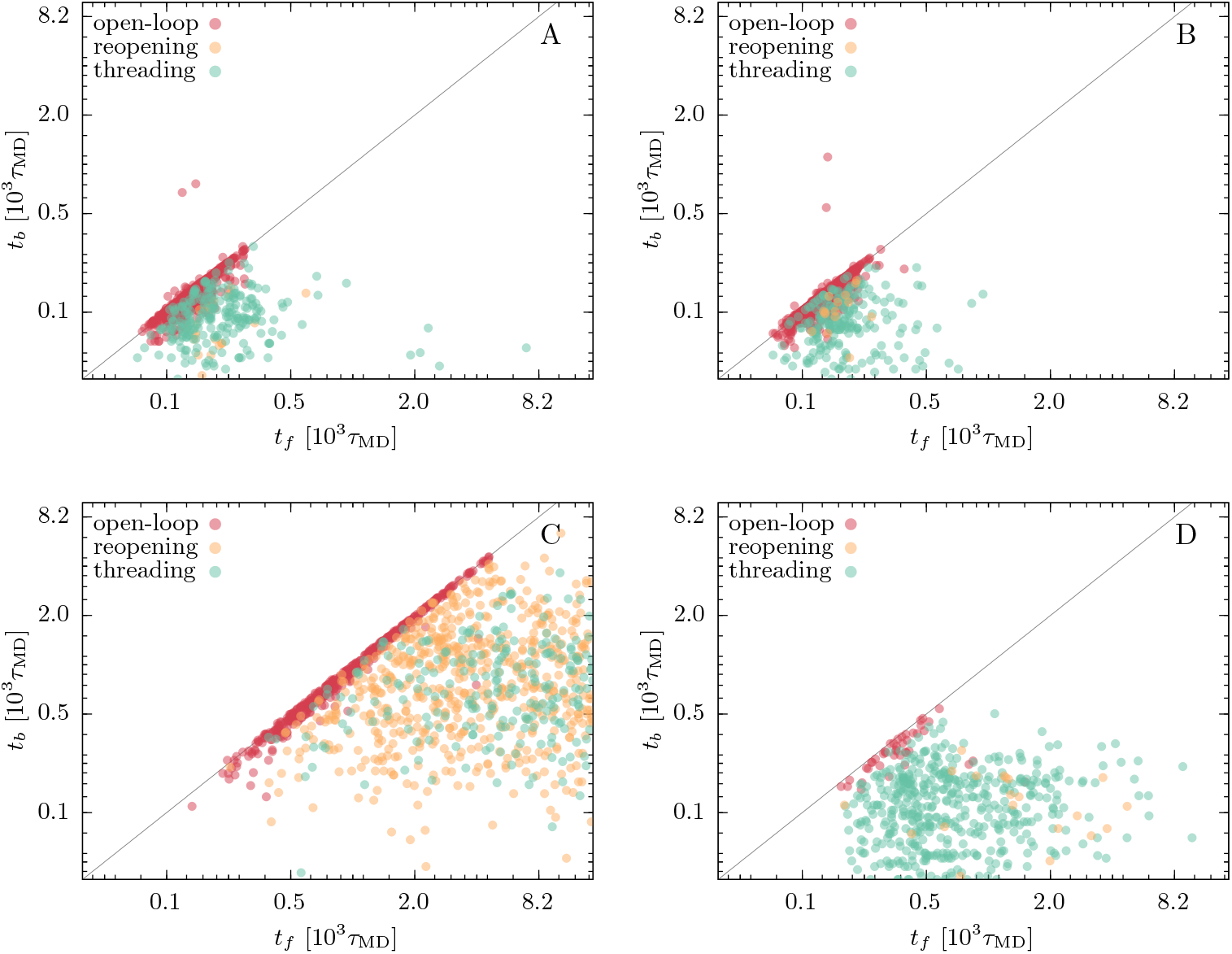
*t_b_* versus *t_f_* for the successful trajectories of GōM simulations. Panels A and B display the results of the GōM at *T* = 0.7: the *N* = 1228 successful trajectories in reducing conditions are shown in panel A and the *N* = 1130 successful trajectories in oxidizing conditions are shown in panel B. Panels C and D display the results of the GōM at *T* = 1.1: the *N* = 1291 successful trajectories in reducing conditions are shown in panel C and the *N* = 549 successful trajectories in oxidizing conditions are shown in panel D. The color of the circles indicates the folding pathway, following the classification indicated in the text. The black line corresponds to *t_b_* = *t_f_*.

Moreover, almost all folding events take place at early times (*t* < 10^3^), while only a minor fraction of trajectories folds in the remaining simulation length. This indicates that the non-successful runs have reached deep, metastable states and would need much longer times to find their way to the native basin. We thus notice that this Gō-like description of 2GMF is prone to kinetic traps, hampering the reproducible folding of the model. Backtracking is here a crucial factor in determining the access to the native state but, at this temperature, it would necessitate much longer timescales than those accessed by our simulations. The optimized EFM model could instead reach a very high probability of folding within the early stages of dynamics. This supports the idea that concerted, non-local motions of the backbone, as those driven by EFM angular potentials, are crucial for reproducible and efficient folding of self-entangled proteins. A model (almost) purely driven by native contacts misses this aspect, and thus fails in folding reproducibly.

To further enrich this picture we show the behavior of the GōM when the folding temperature is increased, facilitating the backtracking mechanism. To this purpose we have studied the GōM at *T* = 1.1, both in reducing and oxidizing conditions. Again, we have collected 2048 runs of length τ_run_ = 1.5 × 10^4^, to detect the fast folding events. As reported in Tab. 1, the probability of folding within this simulation time is now strongly affected by the environment, with a much lower success rate in oxidizing conditions. To investigate the reason for this difference we have collected the distribution of folding times, once again distinguishing among the different pathways. The results, displayed in Fig. 12, show that the process at *T =* 1.1 is on average much slower than at *T* = 0.7, and that the population of folding routes is not symmetric anymore between reduced and oxidized model.

At *T* = 1.1 the model is outside the kinetically optimal regime, and slower routes, that at *T* = 0.7 are prevented by the roughness of the free-energy surface, are made accessible by thermal fluctuations, that allow backtracking and the exploration of the folding funnel across different pathways. This aspect is evident from the behavior of the model in reducing conditions (Fig. 12 C), where the all three mechanisms are well populated, and the incidence of slower pathways is limited only by the finite sampling time of the trajectories. In the oxidized model (Fig. 12 D) the situation is different, since the cysteine bridge potential anticipates the closure of the loop, narrowing the conformational space accessible by thermal fluctuations, and polarizing the choice of folding mechanism towards the threading pathway. The promotion of a single folding route has here a different nature than in EFM results, where it was determined by the optimality of folding kinetics. It would be therefore of great interest to verify the preferential folding pathway of 2GMF by means of more detailed all-atom MD simulations, or with experimental probing. This kind of evidence, on the basis of the results presented here, would indeed provide insights on the nature of folding mechanism selection, that is a characterizing feature of self-entangled proteins.

## CONCLUSION

In this work we have presented an investigation on the folding of the glycoprotein Granulocyte-macrophage colony-stimulating factor (2GMF), which presents a Complex Lasso native structure. The study is performed by means of MD simulations, employing both the widely used Gō Model, proposed by Clementi et al. (26) and the EFM, a Coarse Grained, minimalistic description proposed in Ref. (48) for investigating the folding mechanisms of knotted proteins. We here extended the original models by implementing the formation of native cysteine bridges, in order to assess their effect on the folding process.

The EFM dynamics is based on optimized bending and dihedral potentials, which are tuned to improve the folding capability of the model, with the purpose of enlightening the optimal pathways towards the native structure. In this work we have introduced the MFFO, an evolutionary approach for the optimization of EFM interaction potentials. The results show that this algorithm significantly outperforms the original stochastic method, allowing the study of more complex systems with EFM. Moreover, this evolutionary strategy is general, and can be employed to optimize other minimalistic protein descriptions, such as Gō-like models. Relying on this evolutionary approach, we have built an optimized model of 2GMF, capable of reaching a very high success rate during the early stages of the folding, avoiding kinetic traps, and providing indications on the pathways that enable efficient and reproducible folding. We have then compared the behavior of this model to the results obtained with the GōM.

In our study we focused on the capability of folding in relatively short times, that is, without encountering major kinetic traps. The optimized EFM is in this sense more successful, attaining a folding probability of 0.95 against the 0.6 achieved by the considered Gō-model at the temperature of fastest folding. This demonstrates the importance of force-field heterogeneity, and concerted angular motions, for efficiently crossing the topological bottlenecks of self-entangled folding.

Besides the capability of reaching the native state, we were also interested in studying the folding pathways of the protein. To this purpose we have defined two topological descriptors, the lasso-variable *L*, and the Gauss linking number *G*, inspired by successful methodologies for the classification of protein structures. By monitoring the topology of the protein, these variables turned out to be useful tools for the analysis and classification of folding trajectories. As a result, we were able to characterize the folding scenario of 2GMF, outlining three main mechanisms. Building on this picture, we showed that the optimization of EFM can polarize the trajectories towards an open-loop folding route, in which the lasso topology sets in before the cysteine bridge is formed and seals the covalent loop. The selection of this optimal pathway is also confirmed by the GōM that, at the temperature of fastest folding, privileges an open-loop folding route.

By simulating the GōM at a higher temperature, to lower the free-energy barriers and allow for backtracking mechanism, we have found that the scenario of folding pathways changes. These temperature conditions fall outside the range of optimal folding kinetics, and the process requires much longer simulation times. Nonetheless, as native contacts can break more easily, the protein can sample a larger portion of the free-energy landscape, populating all possible folding routes. Also at this temperature regime, under oxidizing conditions, we have observed a polarization of the folding pathway towards a single mechanism. However, differently from the optimal kinetic scenario, these simulations favor a loop-threading route. Indeed, the early formation of the covalent loop, imposes an entropic restraint to the model, restricting the possible routes to the threading one. Starting from this picture, we think that the study of 2GMF folding using further techniques, either more detailed simulations or experimental studies, would be crucial to validate the hypothesis that entangled folding has evolved to privilege optimal pathways. Overall, this discussion can provide a useful viewpoint in the debate on protein folding mechanisms, and their driving principles (see e.g. (71–73)).

The methodological advancements presented here constitute a useful complement to the existing protein models. They can provide valuable insights on the folding landscape of topologically complex proteins, and draw the guidelines for molecular simulations using more detailed physical models. Moreover, by highlighting the most efficient folding routes, the qualitative picture obtained with the EFM can also shed light on the role played by environmental factors that accelerate folding, such as chaperonins or cotranslational folding.

## AUTHOR CONTRIBUTIONS

C. Perego and R. Potestio designed the research and developed the methodologies. C. Perego carried out all simulations and analyzed the data. C. Perego and R. Potestio wrote the article.

## ACKNOWLEDGMENTS

The authors thank O. Valsson and Y. Zhao for a critical reading of the manuscript. The authors acknowledge funding from the European Union’s Horizon 2020 research and innovation programme under the GOKNOT Marie Skłodowska-Curie Grant Agreement No. 796969, and the contribution of the COST Action CA17139. Computational resources were provided by The Max Planck Computing and Data Facility.

