## Supplemental Material Text for "Searching the optimal folding routes of a Complex Lasso protein"

### S1 Generation of Hybrid Force-fields

In this section we describe the protocol used to generate the coefficients of the hybrid force-fields in the MFFO method. As mentioned in the main submission, after each optimization step, the set of new candidates $\{K'_k\}_{k=1}^{N_K}$ is composed by the winners, namely the $N_{\text{win}}$ that had the best ranking in the previous step, and by $N_K - N_{\text{win}}$ hybrid force-fields. The latter are generated via a crossover operation, or recombination, that mixes the $k_i^{\text{ang}}$ coming from selected "parent" force-fields, mimicking the chromosomal crossover in biology. The parent force-fields are composed by the winners and by $N_{\text{low}}$ low-fit force-fields, introduced to maintain variability in the population. In our calculatons we have generated the low-fit forcefields by randomly picking the $k_i^{\text{ang}}$'s from a uniform distribution ranging between $k_{\text{min}}$ and $k_{\text{max}}$.

Once the parent set is defined the crossover is performed in the following way. As shown in Fig. 12 of the manuscript, crossover points along the backbone are defined, at which the angular coefficients of the parent force-fields are divided in subsets. In our calculations we have defined two crossover points, between residues 42 and 43 and between residues 84 and 85, that is:

$$K = \{k_1^{\text{bend}}, \dots, k_{119}^{\text{bend}}, k_1^{\text{tor}}, \dots, k_{118}^{\text{tor}}\} = \{K_1, K_2, K_3\}, \tag{S1}$$

where the subsets contained both bending and torsion $k$'s:

$$K_1 = \{k_1^{\text{bend}}, \dots, k_{42}^{\text{bend}}, k_1^{\text{tor}}, \dots, k_{42}^{\text{tor}}\}, \tag{S2}$$

$$K_2 = \{k_{43}^{\text{bend}}, \dots, k_{84}^{\text{bend}}, k_{43}^{\text{tor}}, \dots, k_{84}^{\text{tor}}\}, \tag{S3}$$

$$K_3 = \{k_{85}^{\text{bend}}, \dots, k_{119}^{\text{bend}}, k_{85}^{\text{tor}}, \dots, k_{118}^{\text{tor}}\}. \tag{S4}$$

Subsets coming from different parents were recombined at the crossover points to generate the hybrid candidates, formally:

$$H = \{K_1^i, K_2^j, K_3^k\}, \tag{S5}$$

in which $i, j$ and $k$, the indexes of the original parent force-field, were randomly picked among all possible combinations (with no repetition).

### S2 Geometry of Topological Variables

In this section we discuss the structure reduction operated to compute the topological variables defined in the Methods section of the manuscript. We consider the 1-residue-to-1-bead CG representation of

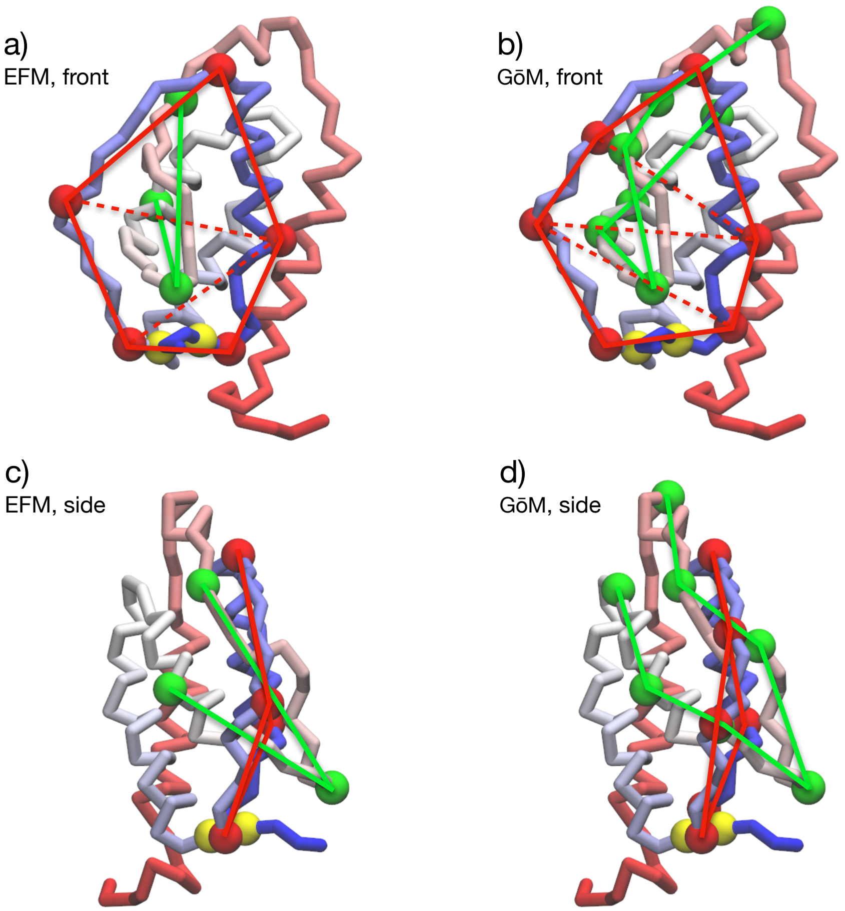


Figure S1: **CG representation of the reduced structures employed for the calculation of $L$ and $G$.** The loop residues $l'$ are highlighted as red circles connected by red lines, while the threading hairpin residues $t'$ are highlighted as green circles connected by green lines. The red dashed lines indicate the triangulation of the loop surface. The structure reduction adoptd for the analysis of EFM trajectories is indicated in a) (front view) and c) (side view), while that employed for the GōM trajectories is indicated in b) (front view) and d) (side view).

the 2GMF protein, adopted in both the EFM and Gō model simulations presented in the manuscript. In this model the protein is described as a polymer chain of 121 monomers, that we label via the index $i = 1, \ldots, 121$ (the first 3 residues are not resolved in the PDB, the real sequence index of the residues is therefore $j = i + 3$). This CG picture of the protein is displayed in Fig. S1.

The first step required to compute the topological variables is to define $l_1, \ldots, l_{N_l}$, namely the indexes of the covalent loop monomers, and $t_1, \ldots, t_{N_t}$, namely the indexes of the threading hairpin monomers. The covalent loop of 2GMF is formed by the $b_1$ cysteine bridge, connecting the monomers 85 and 118, highlighted in figure by yellow spheres, we have thus set $l = 85, \ldots, 118$. In the native fold, the covalent loop is pierced by an hairpin formed by residues from $i = 40$ to $i = 50$. In order to include possible fluctuations of the structure we considered a larger set of residues defined by $t = 30, \ldots, 64$. The next step is the definition of the reduced loop and hairpin indexes, $l'_1, \ldots, l'_{M_l}$ and $t'_1, \ldots, t'_{M_t}$ respectively.

As mentioned in Results section of the manuscript, we have operated two different choices for the reduction, depending if the trajectory was produced via EFM or GōM simulation. In the first case we have represented the loop by residues $l' = 86, 92, 99, 112, 117$ and the hairpin by residues $t' = 37, 47, 57$, as depicted in Figs S1a) and c). The $M_l - 2 = 3$ triangles spanning the loop surface are also indicated in Fig. S1a. These indexes were used for computing the lasso variable $L$, while for the Gauss linking number we also added the cysteine residues 85 and 118 to the definition of the loop.

In the GōM case the dihedral stiffness is on average lower than in EFM ($k_1^{\rm tor} = 1$ and $k_3^{\rm tor} = 0.5$), and the temperature of interest is larger ($T = 0.1$ in EFM runs while $T = 0.7$ or 1.1 in GōM runs). For this reason the secondary structures are less rigid, and we needed to include more monomers in the definition of the topology. We have represented the loop by residues $l' = 86, 91, 95, 99, 112, 116$ and the hairpin by residues $t' = 30, 37, 41, 47, 51, 57, 64$, shown in Figs S1b) and d). The $M_l - 2 = 4$ triangles spanning the loop surface are shown in Fig. S1b.

### S3 Optimized Forcefields

In this section we report the coefficients of the optimized forcefields adopted for the EFM study presented in the main manuscript. In Fig. S2 the bending (A) and torsion (B) stiffnesses for the OM and HM model are displayed. The latter are equal to the average values of the optimized bending and torsion coefficients.

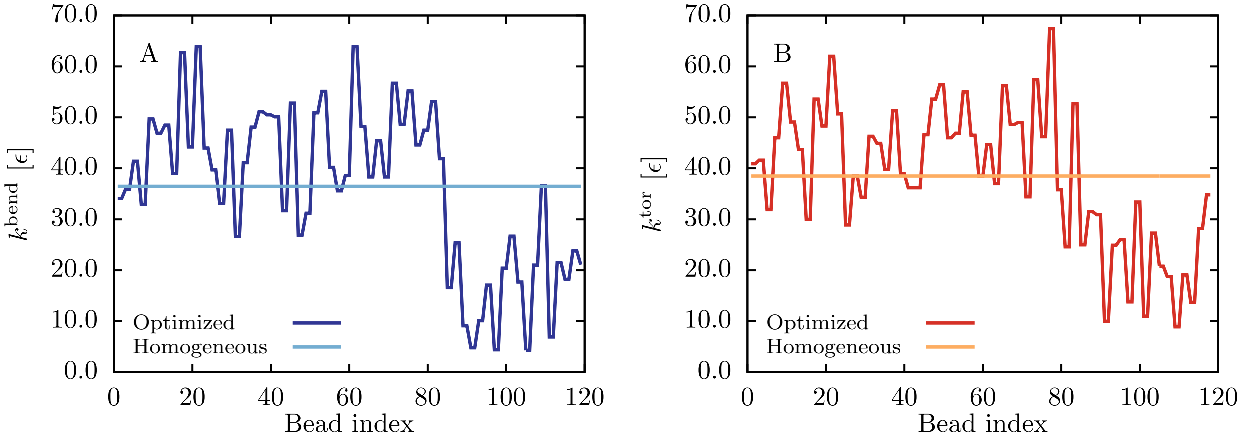


Figure S2: **Angular coefficients of the optimized and homogeneous models, OM and HM**. Panel A and B display the bending and torsion stiffness, respectively.

### S4 Gō Model Temperature Study

The GōM used in the present paper was proposed by Clementi et al. to investigate the folding of small globular proteins[1], as mentioned in the manuscript we have generated the model by means of the SMOG web server (http://smog-server.org)[2,3]. Before comparing the results of the GōM with the EFM simulations we have performed a study on the folding propensity of the GōM at different temperatures, in order to find the range of optimal folding kinetics, at which the fastest folding occurs[4]. We have simulated the folding of 2GMF under reductive conditions, over a range of temperatures from $T = 0.1$ to $T = 1.2$, with spacing $\Delta T = 0.1$. For each value of $T$ we have performed a set of 1024 GōM folding runs of length $\tau = 3500$ and estimated the folding probability $P_f$ and time $t_f$. Our estimate of $P_f$ is equal to the frequency of folding events along the trajectories, it is therefore dependent on the simulation length. Our choice of $\tau$ is justified by the measured median folding times $t_{f,1/2}$, displayed in Fig. S3, where the estimated $P_f$ at different temperatures is displayed as well. Based on these results we have selected to study the GōM model at $T = 0.7$.

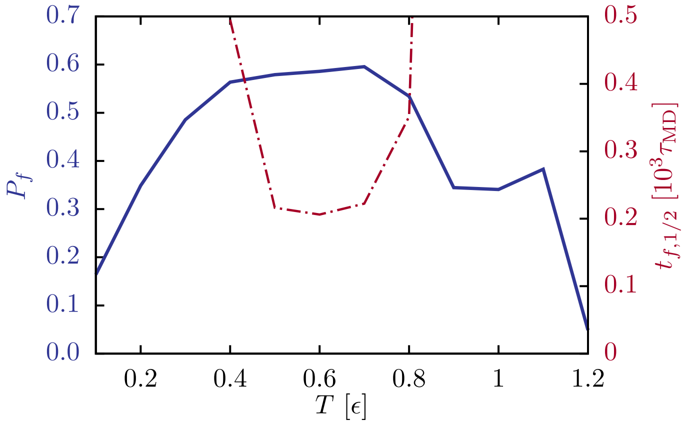


Figure S3: **Temperature range of fastest folding for the GōM.** Folding probability $P_f$ (blue, solid line, left y-axis) and median folding time $t_{f,1/2}$ (red, dot-dashed line, right y-axis) of the GōM at different temperatures.
